# Endogenous α-SYN protein analysis from human brain tissues using single-molecule pull-down assay

**DOI:** 10.1101/188169

**Authors:** Goun Je, Benjamin Croop, Sambuddha Basu, Jialei Tang, Kyu Young Han, Yoon-Seong Kim

## Abstract

Alpha-synuclein (α-SYN) is a central molecule in Parkinson’s disease pathogenesis. Despite several studies, the molecular nature of endogenous α-SYN especially in human brain samples is still not well understood due to the lack of reliable methods and the limited amount of bio-specimens. Here, we introduce α-SYN single-molecule pull-down (α-SYN SiMPull) assay combined with in vivo protein crosslinking to count individual α-SYN protein and assess its native oligomerization states from biological samples including human postmortem brains. This powerful single-molecule assay can be highly useful in diagnostic applications using various specimens for neurodegenerative diseases including Alzheimer’s disease and Parkinson’s disease.

## Introduction

Alpha-synuclein (α-SYN) is a central molecule in Parkinson’s disease (PD) pathogenesis whose aggregates are a major component of Lewy bodies, a pathological hallmark of PD^1^. Both missense point mutations and increased expression of wild-type α-SYN by either multiplication of the SNCA genomic locus or other causes including environmental toxins accelerate α-SYN aggregation and its toxicity^2-6^, suggesting the importance of α-SYN protein levels and oligomeric states in PD pathogenesis.

The characteristics of α-SYN have been mainly studied by using recombinant proteins^7-11^. However, analysis of endogenous α-SYN levels and its aggregation states especially from human brain tissues has been challenging and often reported equivocal results^12-16^. Especially, selective neuronal loss in the substantia nigra (SN) of PD brains severely limits the available dopaminergic neurons compared to control brains^17,18^. Therefore, it is crucial to develop a method capable of quantifying endogenous α-SYN protein levels as well as its aggregation states in biological samples including human postmortem brains.

To this end, we sought to develop a specific and sensitive method called ‘α-SYN single-molecule pull-down (α-SYN SiMPull) assay’ using a recently developed SiMPull method^19,20^ together with cell permeable in vivo crosslinker^21^, disuccinimidyl glutarate (DSG)^13^. DSG has been known to preserve the native state of oligomeric proteins in dynamic equilibrium^22^. A SiMPull method using conventional immunoprecipitation combined with single-molecule fluorescence imaging enables rapid and sensitive analysis of protein levels and their stoichiometry at the single-protein resolution^19-21,23,24^. In addition, in vivo crosslinker increases the stability of endogenous α-SYN protein by preserving their apparent assembly states^13^. Here, we have established α-SYN SiMPull assay, successfully demonstrating that endogenous α-SYN protein levels can be measured at the single-molecule level using only minute amounts of total proteins compared to traditional western blot analysis. Moreover, this method allows us to assess oligomerization states of native α-SYN protein in the cultured cells and the human brain tissues.

## Results and discussion

### Establishing α-SYN SiMPull assay

To achieve specific pull-down of α-SYN protein, we prepared four-antibody system consisting of biotinylated secondary antibody, capturing and detecting primary monoclonal antibodies recognizing different epitopes of α-SYN, and Alexa 647-labeled secondary antibody (Scheme 1)^20^. With this system, recombinant human α-SYN protein was successfully detected by single-molecule fluorescence microscopy (Figure 1a)^25^. Next, we tested whether this method is specific enough to selectively capture α-SYN in total cell lysates. For this purpose, we have established α-SYN knockout 293T cells using the CRISPR/Cas9-based genome editing technique. Then, total lysates prepared from α-SYN knockout and overexpressed cells were each tested. The assay successfully pulled down α-SYN from α-SYN overexpressed cell lysates with high specificity when compared to negligible signals from α-SYN knockout cell lysates (Figure 1b,d). We also applied α-SYN SiMPull to detect endogenous α-SYN from total lysates of wild-type 293T cells, demonstrating that the number of fluorescence spots was increased in a dose-dependent manner (Figure 1c,d).

**Scheme 1.**
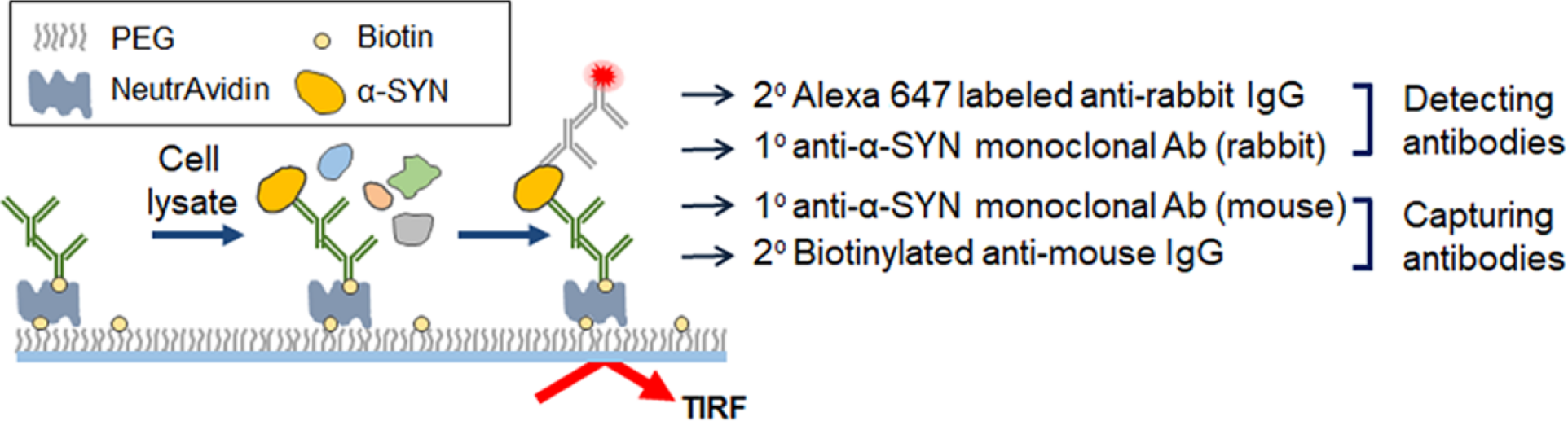
Schematic diagram of α-SYN SiMPull procedure with four-antibody system.

**Figure 1.**
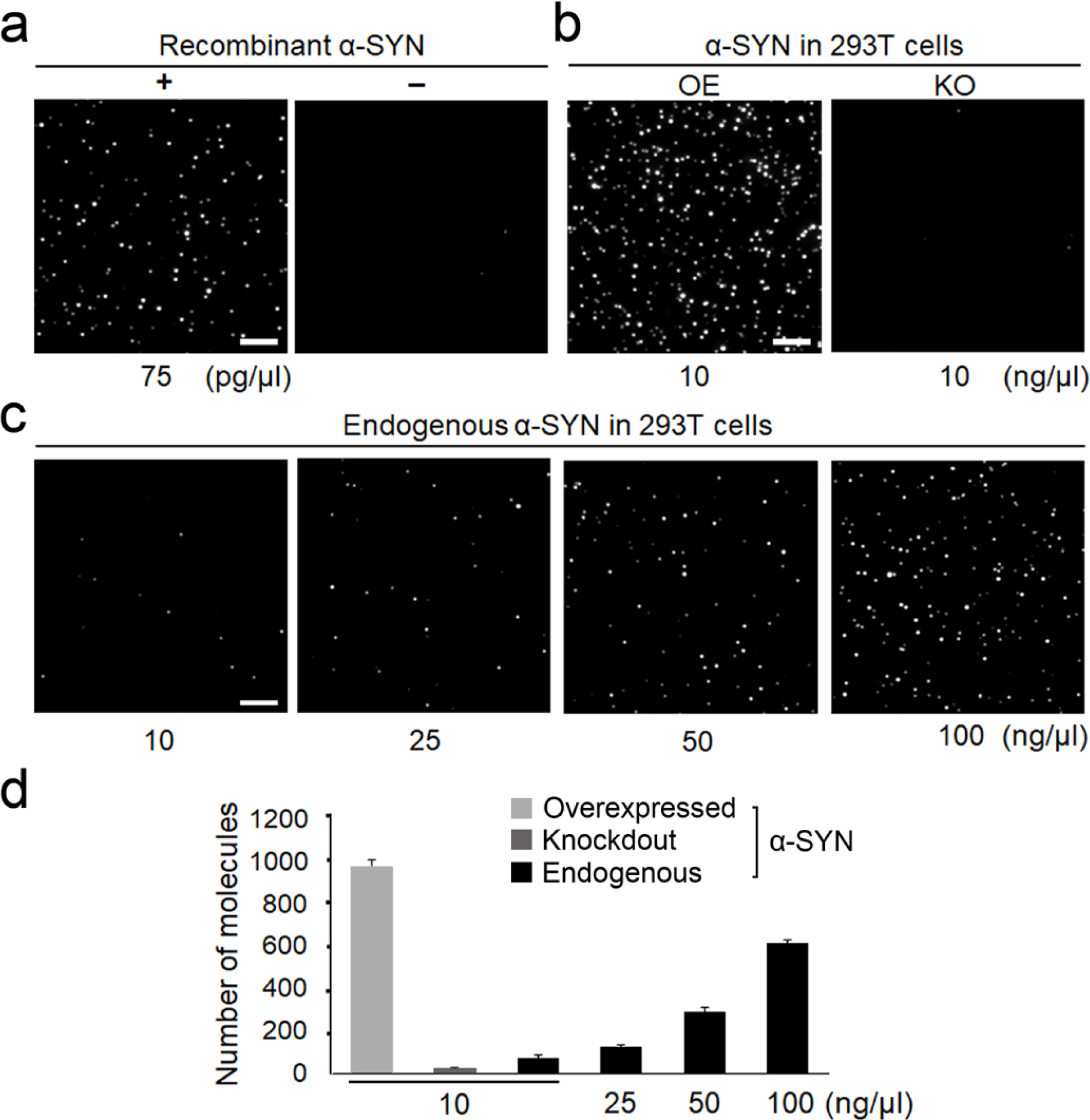
Detection of α-SYN protein using SiMPull assay. (a) Single-molecule images of recombinant human α-SYN protein (75 pg/μL) (left) and non-specific binding of Alexa 647-labeled anti-rabbit IgG (right). (b) Single-molecule images of α-SYN taken from total lysates of α-SYN overexpressed (OE) or knockout (KO) 293T cells with 10 ng/μL of total lysates. (c) Images of endogenous α-SYN from 293T cells with 10, 25, 50, or 100 ng/μL of total lysates. (d) Average number of fluorescent spots of α-SYN molecules per imaging area. More than 20 images were taken and error bars denote standard deviation (s.d.). Scale bar, 5 μm. All data are representatives of three independent experiments.

### Analysis of oligomeric states of recombinant α-SYN by α-SYN SiMPull assay

α-SYN oligomerization is strongly implicated in mediating α-SYN toxicity in neurons^26,27^. Therefore, understanding the oligomerization states of α-SYN is important for diagnosing as well as monitoring the progression of PD. To study α-SYN oligomers, first, we adopted Alexa 647-labeled F(ab’)_2_ fragment antibody instead of full IgG to reduce the steric hindrance between antibodies, and used a degree of labeling of ˜2.9 to achieve a narrow fluorescence intensity distribution with a negligible amount of unlabeled species (Figure S1). Then SiMPull assay was performed on human α-SYN oligomer prepared by 5-day incubation of recombinant monomer at 37 degrees (Figure 2a-e)^28^. We analyzed the fluorescence intensity of the immuno-precipitated molecules, which is proportional to the number of α-SYN in the pulled-down complexes. As we expected, in the oligomeric/fibrillar α-SYN sample, multiple bright spots with various intensity were observed while the monomeric α-SYN sample depicted a narrow distribution centered at low intensity (Figure 2d,e). Interestingly, we also observed fluorescent spots with various shapes, which are larger than the diffraction limit (˜ 350 nm) exclusively in oligomeric/fibrillar α-SYN (Figure 2b,c), suggesting that α-SYN SiMPull assay is applicable to morphometric analysis. Monomeric and oligomeric α-SYN were confirmed using conventional western blot with significantly higher amount of proteins (Figure S2).

**Figure 2.**
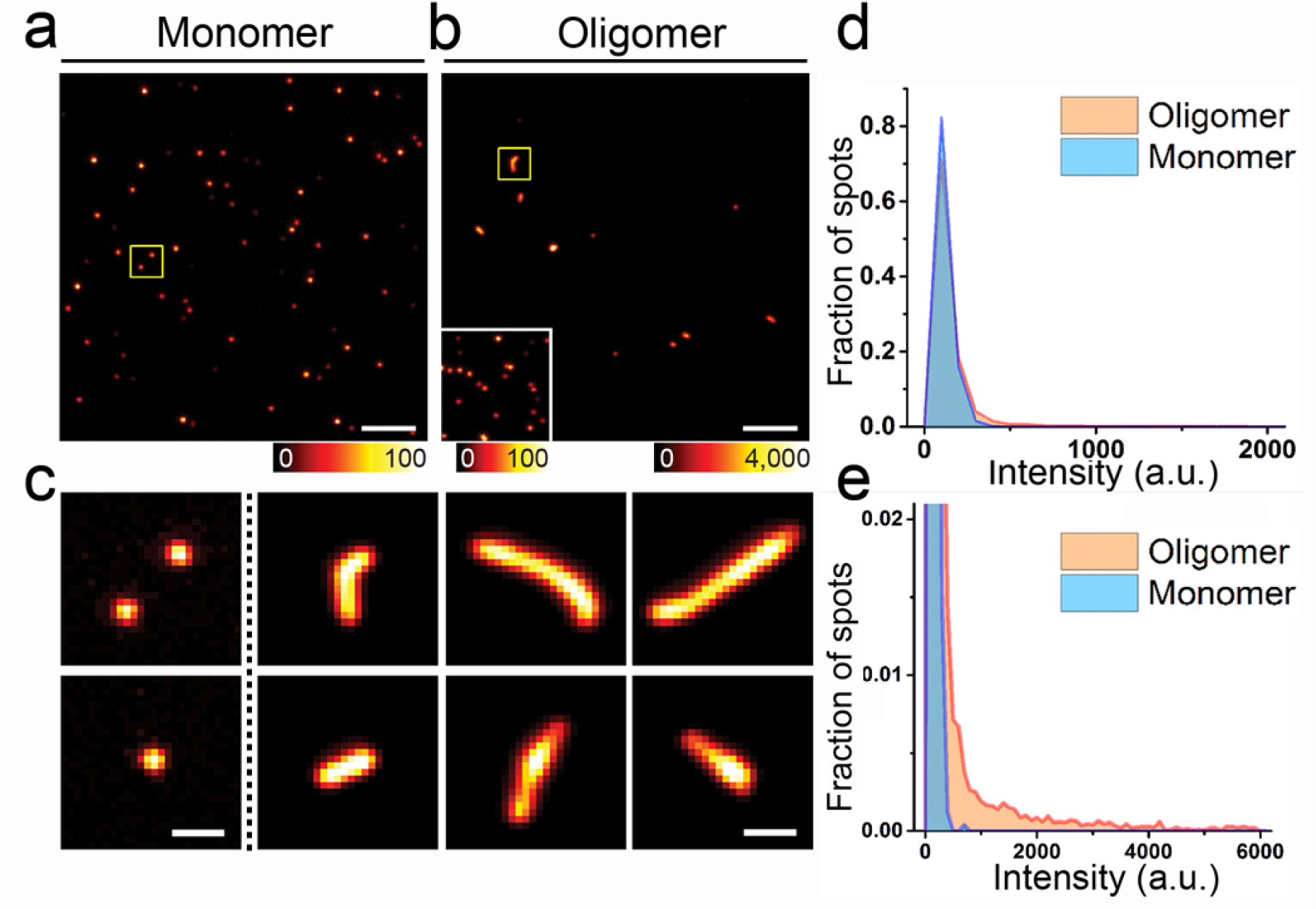
Analysis of oligomeric states of recombinant α-SYN using SiMPull assay. Single-molecule images of monomeric (a) and oligomeric (b) recombinant α-SYN (37.5 pg/μL). Existing monomers among oligomeric α-SYN were displayed on the left corner of (b) after intensity adjustment. (c) Shapes of monomeric (left-most panel) and oligomeric/fibrillar α-SYN (three right panels). The first two in the upper panel are magnified images of the yellow-boxed area in (a) and (b), and the others were taken from other images. (d,e) Fluorescence intensity profiles of monomeric (blue) and oligomeric (red) recombinant α-SYN. High intensity spots were observed exclusively in oligomeric recombinant α-SYN. Scale bar, 5 μm (a,b) and 1 μm (c). All data are representatives of three independent experiments.

### Analysis of α-SYN in the cultured cells by α-SYN SiMPull assay

To extend the application of α-SYN SiMPull to analysis of α-SYN oligomeric states in the cells, we tested total lysates from 293T cells overexpressing α-SYN with or without exposure to FeCl_2_ and a proteasome inhibitor, MG132, which are known to increase aggregation of α-SYN^29-31^. To stably maintain the native states of α-SYN, in vivo protein crosslinking was achieved by DSG followed by cell lysis^13^. α-SYN SiMPull using 40 μL of total lysates (10 ng/μL) demonstrated notably increased population at higher fluorescence intensity in FeCl_2_ and MG132 treated cells compared to non-treated control (Figure 3a and Figure S3). To further analyze these intensity profiles, fluorescence intensity of a single Alexa 647-labeled F(ab’)_2_ was used as a reference (Figure S1a,b). Assuming it as the intensity of a monomer, we decomposed the intensity profiles of above conditions into monomer and oligomer populations. As shown in Figure 3b and c, oligomeric α-SYN was more pronounced in FeCl_2_ and MG132 exposed cells (37 %) than non-exposed ones (15 %). The western blot also confirmed that the treatment with FeCl_2_ and MG132 increased the levels of high-molecular weight α-SYN species (Figure S4). However, we did not observe bright fluorescent spots having different shapes or sizes that were found in recombinant oligomeric/fibrillar α-SYN in FeCl_2_ and MG132 exposed cells.

**Figure 3.**
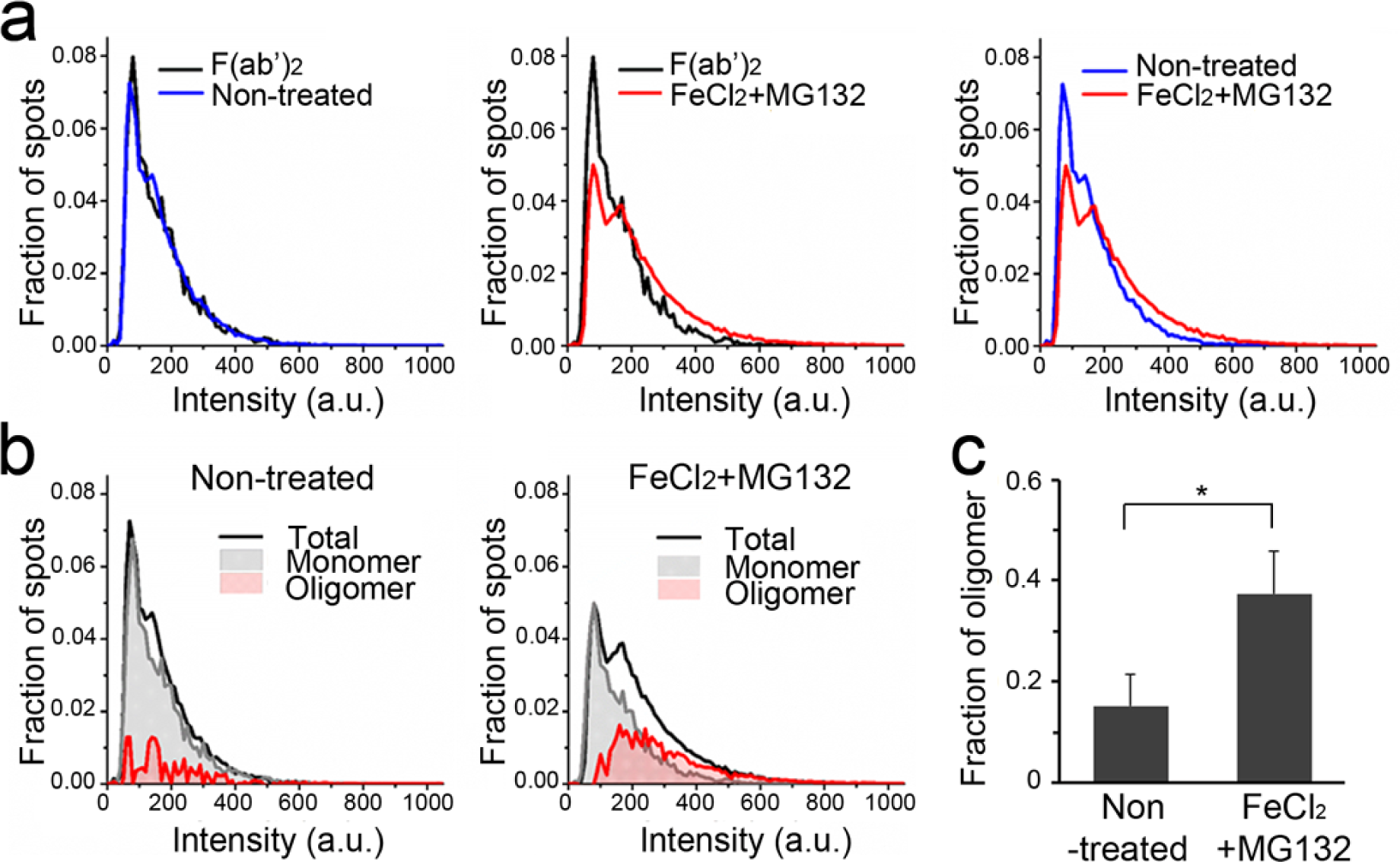
Analysis of oligomeric states of α-SYN from in vivo DSG-crosslinked total cell lysates using SiMPull. (a) Fluorescence intensity distribution of α-SYN SiMPull assay from α-SYN overexpressed cells with (red) or without (blue) FeCl_2_ and MG132 treatment. F(ab’)_2_ denotes Alexa 647-labeled F(ab’)_2_ fragment antibody as a reference of α-SYN monomer. (b) Analysis of oligomeric states from the data presented in (a). Monomeric (grey) and oligomeric (red) populations were separately plotted. (c) Quantitative analysis of the oligomeric states from the data presented in (b). Error bars denote standard error of the mean (n = 3). *P< 0.05, by unpaired two-tailed t test. 10 ng/μL of total lysates from in vivo DSG-crosslinked 293T cells were used in each assay. All data are representatives of three independent experiments.

### Analysis of α-SYN in the human brain tissues by α-SYN SiMPull assay

Lastly, we applied α-SYN SiMPull assay to test human postmortem brain samples. The dark pigmented region in the SN of frozen control or PD postmortem brain samples that represents remaining dopaminergic neurons was selectively punch-biopsied (˜10 mg) to minimize compounding effects contributed by other cells, and then treated with DSG for in vivo crosslinking prior to protein extraction (Figure 4a). PD sample showed significant increase in the number of fluorescent spots by 3.3 fold compared to control (Figure 4b). Moreover, the population of oligomeric α-SYN was significantly increased in PD, where it accounted for 56 % of the detected protein, compared to only 23% in control (Figure 4c-e). Neither samples showed different shapes of fibrillar α-SYN. The western blot using significantly higher amount of lysates than SiMPull assay showed prominent levels of high molecular species in PD (Figure S5).

**Figure 4.**
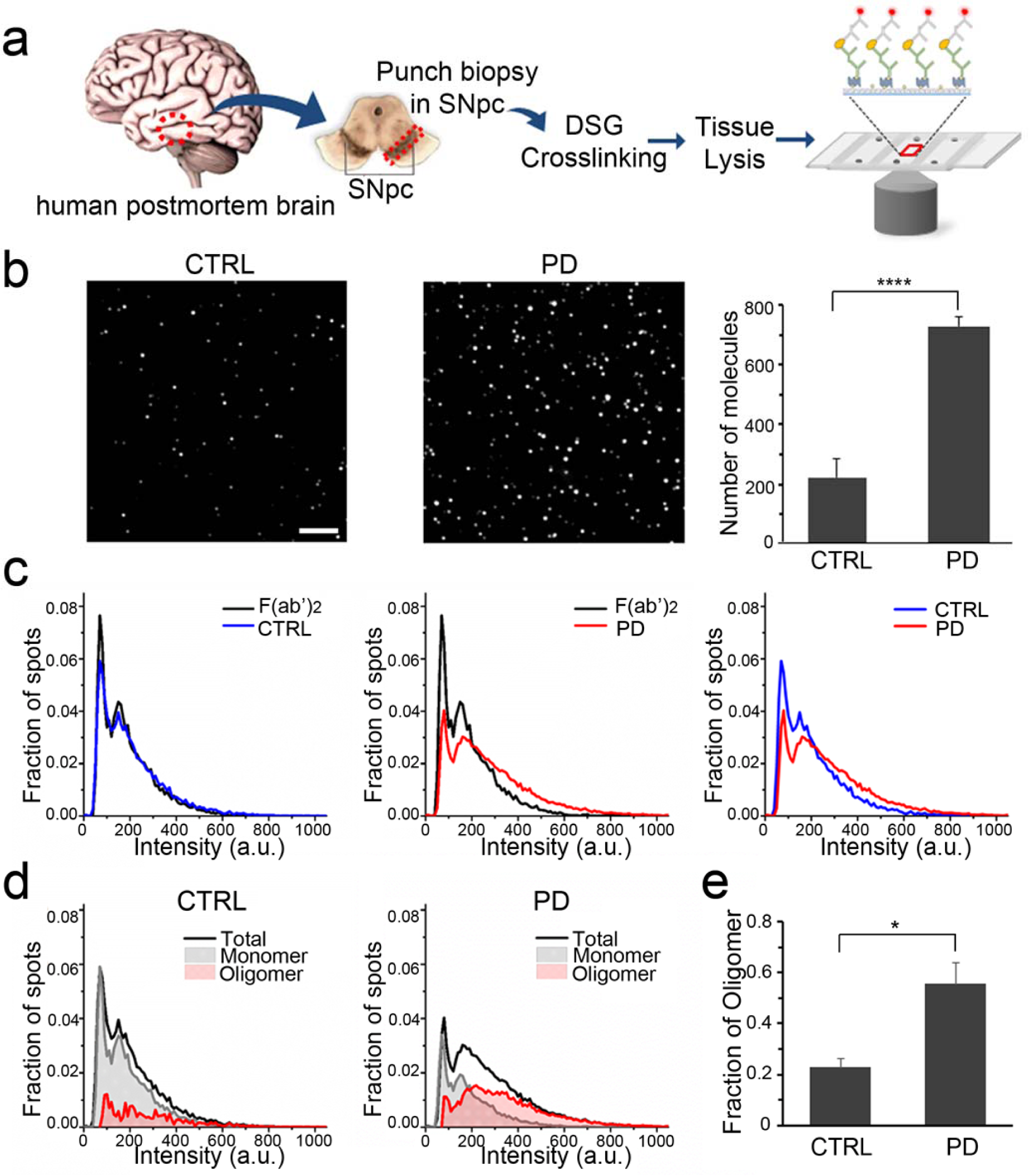
Analysis of oligomeric states of α-SYN from in vivo DSG-crosslinked human postmortem brain punch biopsy samples using SiMPull. (a) Schematic diagram of α-SYN SiMPull procedure from human postmortem brain samples. (b) Single-molecule images of α-SYN from control (CTRL, left) or PD brain samples (middle), and average number of molecules per imaging area (right) taken from 20 images. Scale bar, 5 μm. (c) Fluorescence intensity distribution of α-SYN SiMPull assay from CTRL (blue) and PD (red) brain samples plotted with reference intensity profile of F(ab’)2 (black). (d) Analysis of oligomeric states of CTRL and PD brain samples. Monomeric (grey) and oligomeric (red) populations were separately plotted. (e) Quantitative analysis of the oligomeric states from the data presented in (d). Error bars denote standard deviation (s.d.) in (b) and standard error of the mean (n = 3) in (e). *P< 0.05, ****P< 0.0001 by unpaired two-tailed t test. 50 ng/μL of total lysates from in vivo DSG-crosslinked human control or PD postmortem brain samples were used. All data are representatives of three independent experiments.

## Conclusion

A limited number of remaining dopaminergic neurons in the SN of PD brains have been a critical barrier for biochemical analyses of α-SYN in postmortem PD brain tissues. Several methods including, but not limited to, western blot have reported controversial results on α-SYN levels as well as its native states with total protein lysates obtained from the SN of human brain tissues^12-16^. In this study, we employed α-SYN SiMPull assay that enables us to measure endogenous α-SYN protein at the single-molecule level and estimate its oligomeric states using minute amounts of protein lysates. This technique allowed us to analyze α-SYN in the limited SN region where neuromelanin-positive dopaminergic neurons were spared.

Unlike other methods^15,32,33^, α-SYN SiMPull assay can be applied regardless of the size or conformation of α-SYN oligomers unless the epitope is concealed. We observed fluorescent spots with various shapes in recombinant oligomeric/fibrillar α-SYN, suggesting that different levels of α-SYN aggregation could be analyzed by morphometric study. However, neither cells nor brain samples gave a similar result. It is possible that the effort to maintain the native states of α-SYN by in vivo crosslinking and mild protein extraction procedure is still causing disruption of α-SYN aggregates. Another possibility is that small oligomers are preferably formed while fibrillar α-SYN rarely exists in dopaminergic neurons even in PD. Super-resolution imaging techniques^34^ may extend our method to reveal ultrastructure of the oligomers from PD brain. In the cultured cell experiments, we used FeCl_2_ and the proteasome inhibitor to facilitate aggregation of α-SYN. Alternatively, it is worth to apply this technique to assess seeded aggregation of α-SYN achieved by directly introducing α-SYN preformed fibrils into cells^35^. We observed higher amounts of oligomers in PD brain sample compared to control subject. In this study, we used in vivo crosslinking to stabilize α-SYN oligomers. The potential artifacts mimicking oligomeric states of α-SYN caused by crosslinker were ruled out by demonstrating that control 293T cells expressing α-SYN crosslinked with DSG exhibited very similar intensity profile to Alexa 647-labeled F(ab’)_2_ alone.

Here, we assessed monomeric versus oligomeric α-SYN states using a known fluorescence intensity profile of the Alexa 647-labeled F(ab’)_2_. However, for precise stoichiometric analysis of the native states of α-SYN, a quantitative labeling of the primary antibody^36^ is desirable along with photobleaching analysis^21^. This may provide a crucial clue to solve current debate over the oligomeric states of endogenous α-SYN^12-14^. Adopting smaller antibodies such as Fab fragment or single-chain variable fragment might be more advantageous to bind each molecule in α-SYN oligomers. Further, it is possible to observe other molecules in complex with α-SYN oligomers by multi-color SiMPull assay^20^ and reduce protein amounts by imaging a larger area or constructing microfluidic chambers^37^. Additionally, α-SYN analysis in a single-dopamine neuron could be achieved by in situ single cell pull-down assay^38^. Note that our method can be carried out using any commercial total internal reflection fluorescence (TIRF) microscope system without specialized hardware.

In summary, our α-SYN SiMPull assay will be a powerful tool in quantitation as well as analysis of oligomeric states of various proteins, such as amyloid beta, α-SYN and tau from human postmortem brain tissues or cerebrospinal fluids in neurodegenerative diseases including Alzheimer’s disease and PD.

## Author contributions

G.J., K.Y.H. and Y.S.K. conceived the project. G.J. designed and performed the experiments, and analyzed data. B.C. performed experiments and analyzed data. S.B. prepared α-SYN knockout 293T cells. J.T. built single-molecule TIRF microscope and made MATLAB scripts. G.J., B.C., K.Y.H. and Y.S.K. wrote the manuscript.

## Notes

The authors declare no competing financial interests.

## Acknowledgment

We thank Vasudha Aggarwal and Ankur Jain for critically reading manuscript and fruitful discussions. This work was supported by National Institutes of Health grant number 5R21NS088923 to Y.S.K. and ORC Mentoring Program Award at University of Central Florida to K.Y.H and Y.S.K.

